# Adventitious viruses persistently infect three commonly used mosquito cell lines

**DOI:** 10.1101/317628

**Authors:** James Weger-Lucarelli, Claudia Rückert, Nathan D. Grubaugh, Michael J. Misencik, Philip M. Armstrong, Mark D. Stenglein, Gregory D. Ebel, Doug E. Brackney

**Author notes:** these authors contributed equally. Present address: Department of Biomedical Sciences and Pathobiology, Virginia Polytechnic Institute and State University 360 W Campus Drive, Blacksburg, Virginia, USA. Present address: Department of Immunology and Microbial Science, The Scripps Research Institute, La Jolla, CA, USA.

## Abstract

Mosquito cell lines were first established in the 1960’s and have been used extensively in research to isolate and propagate arthropod-borne (arbo-) viruses, study the invertebrate immune system, and understand virus-vector interactions. Despite their utility as an *in vitro* tool, these cell lines are poorly defined and may harbor insect-specific viruses that could impact experimental results. Accordingly, we screened four commonly-used mosquito cell lines, C6/36 and U4.4 cells from *Aedes albopictus*, Aag2 cells from *Aedes aegypti*, and Hsu cells from *Culex quinquefasciatus*, for the presence of adventitious viruses. All four cell lines stained positive for double-stranded RNA by immunofluorescence, indicative of RNA virus replication. We subsequently identified viruses infecting Aag2, U4.4 and Hsu cell lines using untargeted next-generation sequencing, but not C6/36 cells. Sequences from viruses in the families *Birnaviridae*, *Bunyaviridae, Flaviviridae,* and *Rhabdoviridae* were abundant in the mosquito cell lines. PCR confirmation revealed that these sequences stem from active viral replication and/or integration into the cellular genome. Our results show that these commonly-used mosquito cell lines are persistently-infected with several viruses. This finding may be critical to interpreting data generated in these systems.

## Introduction

Cell culture systems have revolutionized biomedical science and provided key insights into the fundamentals of life. The tractability of these systems make it possible to perform high-throughput drug screens and gene studies (Broach and Thorner, 1996; Perrimon and Mathey-Prevot, 2007), isolate and amplify viruses and develop vaccines (Enders et al., 1949; Lloyd et al., 1936; Rivers and Ward, 1935); experiments that would otherwise be too difficult or impossible to perform *in vivo*. Despite their utility, it has recently been shown that many commonly used mammalian cell lines are persistently infected with a myriad of viruses, possibly confounding the results generated in these cell lineages and highlighting the need for a better understanding of cell culture systems (Hué et al., 2010; Platt et al., 2009; Uphoff et al., 2010).

Developed in the 1960s (Grace, 1966; Peleg, 1968; Singh, 1967), mosquito cell culture systems have become an indispensable tool in the study of arthropod-borne (arbo)viruses. These systems have provided insights into virus evolution and virus-vector interactions and democratized research by allowing laboratories lacking mosquito facilities to investigate arboviruses (Vasilakis et al., 2009; Walker et al., 2014). In addition, they are routinely used to isolate and amplify arboviruses, specifically *Aedes albopictus*-derived C6/36 cells which are deficient in the primary antiviral pathway, RNA interference (Brackney et al., 2010). These systems are generated by macerating whole mosquito larvae or tissues and culturing amenable cells (Walker et al., 2014). This can be problematic because the culture may be composed of one or more unknown cell types. In addition, environmental contaminants such as insect-specific viruses (ISVs) may be unknowingly co-cultured as has been reported for *Drosophila* and tick cell lines (Bell-Sakyi and Attoui, 2013, 2016; Wu et al., 2010). In fact, ISVs have been identified in many mosquito species and both cell-fusing agent virus (CFAV; *Flaviviridae*) and Phasi-charoen like virus (PCLV; *Bunyaviridae*) have been identified in *Aedes aegypti* Aag2 cells (Maringer et al., 2017; Roundy et al., 2017; Schultz et al., 2018; Stollar and Thomas, 1975). Together these data suggest that commonly used mosquito cells may be persistently infected with unknown viruses and defining the culture virome will be critical to properly interpreting data generated in these systems.

In this study, we investigated the possibility that commonly used mosquito cell lines may be persistently infected with ISVs. Using an anti-dsRNA specific antibody, we performed immunofluorescence on uninfected cultures of Aag2 (*Ae. aegypti*), U4.4 (*Ae. albopictus*), C6/36 (*Ae. albopictus*) and Hsu (*Culex quinquefasciatus*) cells. We observed the presence of dsRNA in each cell line indicating the presence of ostensibly viral RNA. Subsequently, we sequenced RNA from these cell lines by next-generation sequencing (NGS) in order to better characterize the origins of this signal. We taxonomically categorized non-host sequences (Fauver et al., 2016a) to identify full-length or partial viral sequences in all cell lines. We further detected viral RNA by RT-PCR in cell supernatant and/or cell lysates and in some instances, DNA forms of RNA viruses. Together, these data demonstrate that many commonly used mosquito cell culture systems are persistently infected with ISVs; results which should be considered when interpreting data generated in these cell lines.

## Materials and Methods

### Cell lines

The *Cx. quinquefasciatus* ovary-derived Hsu (Hsu et al., 1970), *Ae. albopictus*-derived C6/36 (Singh, 1967), and *Ae. aegypti*-derived Aag2 (Lan and Fallon, 1990; Peleg, 1968) cell lines were maintained at 28°C with 5% CO_2_ in MEM supplemented with 10% fetal bovine serum (FBS), 1X nonessential amino acids (100x; ThermoFisher Scientific), 1% L-glutamine, 1% 100x antibiotic-antimycotic (10,000 mg/ml of streptomycin, 10,000 U/ml penicillin, and 25 mg/ml of amphotericin B), and 5% of a 7.5% sodium bicarbonate solution. *Ae. aegypti*-derived U4.4 cells were maintained at 28°C with 5% CO_2_ in Mitsuhashi and Maramorosch insect medium supplemented with 7% FBS, 1X nonessential amino acids, L-glutamine, and antibiotics-antimycotics (10,000 mg/ml of streptomycin, 10,000 U/ml penicillin, and 25 mg/ml of amphotericin B). RNA was sequenced from three batches of C6/36 cells (two from Colorado State University and one from the Connecticut Agricultural Experiment Station) in order to provide insight into inter-laboratory variability. All three batches were originally acquired from ATCC.

### West Nile virus infections

Mosquito cells were plated in 12-well plates at concentrations between 8.1 × 10^5^ and 1.8 × 10^6^ cells/ well on poly-L-lysine treated coverslips. Cells were infected with West Nile virus (WNV) strain 10679-06 at a multiplicity of infection (MOI) of 0.1. Mock infected cells were treated with media. The inoculated plates were incubated at 28°C for 1 hour, with manual rocking at 15 minute intervals, to allow for virus adsorption. After the incubation period, 1 mL of media was added to each well and plates were placed in a 28°C incubator with 5% CO_2_. Both the experimental and control cells were harvested either 24 or 72 hours post infection (h.p.i.).

### Immunofluorescence

Cells were fixed in well with 4% paraformaldehyde for 20 min. at room temperature. Subsequently, cells were permeabilized (PBS + 0.3% TritonX100) for 10 min. at room temperature and incubated with blocking buffer (5% BSA + 0.1% TritonX100) at 4°C overnight. Coverslips were placed in a humid chamber, 50 μL of primary anti-dsRNA antibody (J2) diluted 1:200 in blocking buffer was added to each, and incubated at room temperature for 1 hour. Coverslips were washed three times in wash buffer (PBS+0.1% Tween 20) and incubated with 50 μL of secondary antibody (Alexa-Fluor 555 α-mouse) in the dark for 1 hour at room temperature. Coverslips were washed three additional times in wash buffer and mounted on glass slides with Prolong Gold anti-fade with DAPI counterstain. Slides were visualized on a Leica SP5 confocal microscope using the 405 nm laser (DAPI; nuclei) and 561 Argon laser (Alexa-Fluor 555; dsRNA) at 63x magnification. Brightness and contrast from resultant images were adjusted manually in Adobe Illustrator. All images were adjusted equally.

### Next-generation sequencing of cellular RNA

RNA from cell lines was extracted using the Qiagen viral RNA kit and prepared for sequencing as previously described (Grubaugh et al., 2016). Briefly, each sample was DNase treated using Turbo DNase (Ambion). Total RNA was then non-specifically amplified and converted into dsDNA using the NuGEN Ovation RNA-Seq System V2. dsDNA was then sheared using the Covaris S2 Focused-ultrasonicator according to the manufacturer’s recommendations. Sequencing libraries were prepared from sheared cDNA using NuGEN’s Ovation Ultralow Library Kit according to the manufacturer’s recommendations. Agencourt RNAclean XP beads (Beckman Coulter Genomics, Pasadena, CA) were used for all purification steps. Finished libraries were analyzed for correct size distribution using the Agilent Bioanalyzer High Sensitivity DNA chips (Agilent). 100 nt paired-end reads were generated using the Illumina HiSeq 2500 platform at Beckman Coulter Genomics.

### Virus discovery pipeline

An in-house virus discovery pipeline was used to identify novel viral sequences as previously described (Fauver et al., 2016a). Briefly, reads were first trimmed with cutadapt version 1.13 (Martin, 2011) and then PCR duplicates were removed with CD-HIT-EST tool, version 4.6 (Li and Godzik, 2006). Sequences that mapped to the *Ae. aegypti* (GCF_002204515.2), *Ae. albopictus* (GCF_001876365.2), *An. Gambiae* (GCF_000005575.2), or *Cu. Quinquefasciatus* (GCF_000209185.1) genome assemblies were then removed by alignment with Bowtie2 (Langmead and Salzberg, 2012). Remaining reads were assembled using the SPAdes genome assembler (Bankevich et al., 2012). The contigs produced were then aligned to the NCBI nucleotide database using BLASTn (Altschul et al., 1997; Camacho et al., 2009). Contigs that did not align at the nucleotide level with an e-value less than 10^−8^ were then used for a translation-based search against protein sequences using the DIAMOND (Buchfink et al., 2015). Contigs whose highest scoring alignments were to virus sequences were manually inspected in Geneious v11 (Kearse et al., 2012), and validated by mapping reads back to assemblies using Bowtie2 as above.

### Viral RNA/ DNA detection by PCR

Approximately 8×10^6^ cells of each cell line (Aag2, C6/36, Hsu and U4.4) were harvested by scraping, equally divided into two separate tubes (one for RNA and one for DNA), and pelleted at 10,000xg at 4C for 5 minutes. Cell supernatant was removed and placed in two separate tubes. DNA was extracted from cell pellets using the Zymo Quick gDNA mini-prep. Samples for RNA extraction were all treated with DNase (Promega, Madison, WI) prior to extraction to remove cellular DNA. One of the tubes of cell supernatant was also subjected to RNase A (Thermofisher, 100μg/mL at 37C for one hour) treatment to remove unencapsidated RNA. RNA was extracted from cell pellets, cell supernatant, and RNase A treated RNA using the Zymo DirectZol RNA extraction kit. cDNA was produced from extracted RNA using Protoscript II RT (NEB) using random hexamers. DNA or cDNA was then used for PCR or qPCR using OneTaq DNA polymerase (NEB) or iTaq SYBR green (Biorad), respectively. All qPCRs were confirmed by running a gel to confirm the result visually. Primers used in this study are listed in Table 1.

**Table 1:**
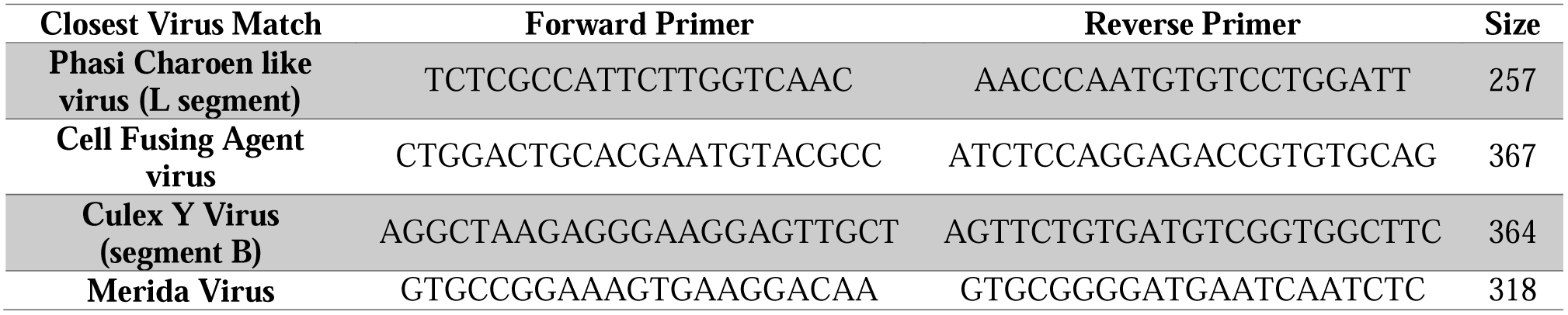
List of primers.

### Data Availability Statement

Raw sequencing reads can be found on the NCBI SRA database (BioProject # PRJNA464394). The accession numbers for the assembled viral contigs are PCLV L segment (MH310079), PCLV M segment (MH310080), PCLV S segment (MH310081), CFAV (MH310082), MERDV (MH310083), CYV segment A (MH310084), and CYV segment B (MH310085). Aliquots of cell lines are available upon request.

## Results

### Detection of dsRNA in mosquito cells

There are numerous reports of invertebrate cell cultures systems persistently infected with ISVs (Bell-Sakyi and Attoui, 2013, 2016; Wu et al., 2010); however, the extent to which commonly used mosquito cell culture systems are infected remains unknown. Therefore, we screened Aag2, C6/36, U4.4, and Hsu cells for the presence of dsRNA. Not normally expressed in eukaryotic cells, dsRNA can be readily detected in cells infected with ssRNA, dsRNA, and DNA viruses using the anti-dsRNA antibody, J2 (Weber et al., 2006). We used WNV infected (24 and 72 h.p.i.) and uninfected cultures of each of the four cell lines as controls for our immunofluorescence assays. As expected, we detected WNV dsRNA in all four cell lines with increasing signal intensity with time post infection suggesting active WNV replication (Fig. 1). Interestingly, we found that all of the uninfected cultures also stained positive for dsRNA (Fig. 1). While Aag2 cells are known to be infected with CFAV (Scott et al., 2010; Stollar and Thomas, 1975) and PCLV (Maringer et al., 2017), and stained positive for dsRNA (Fig. 1A), we also found that a large proportion of C6/36, U4.4 and Hsu cells stained positive for dsRNA (Fig. 1B-D). These data suggest that all four cell lines are persistently infected with at least one virus.

**Figure 1:**
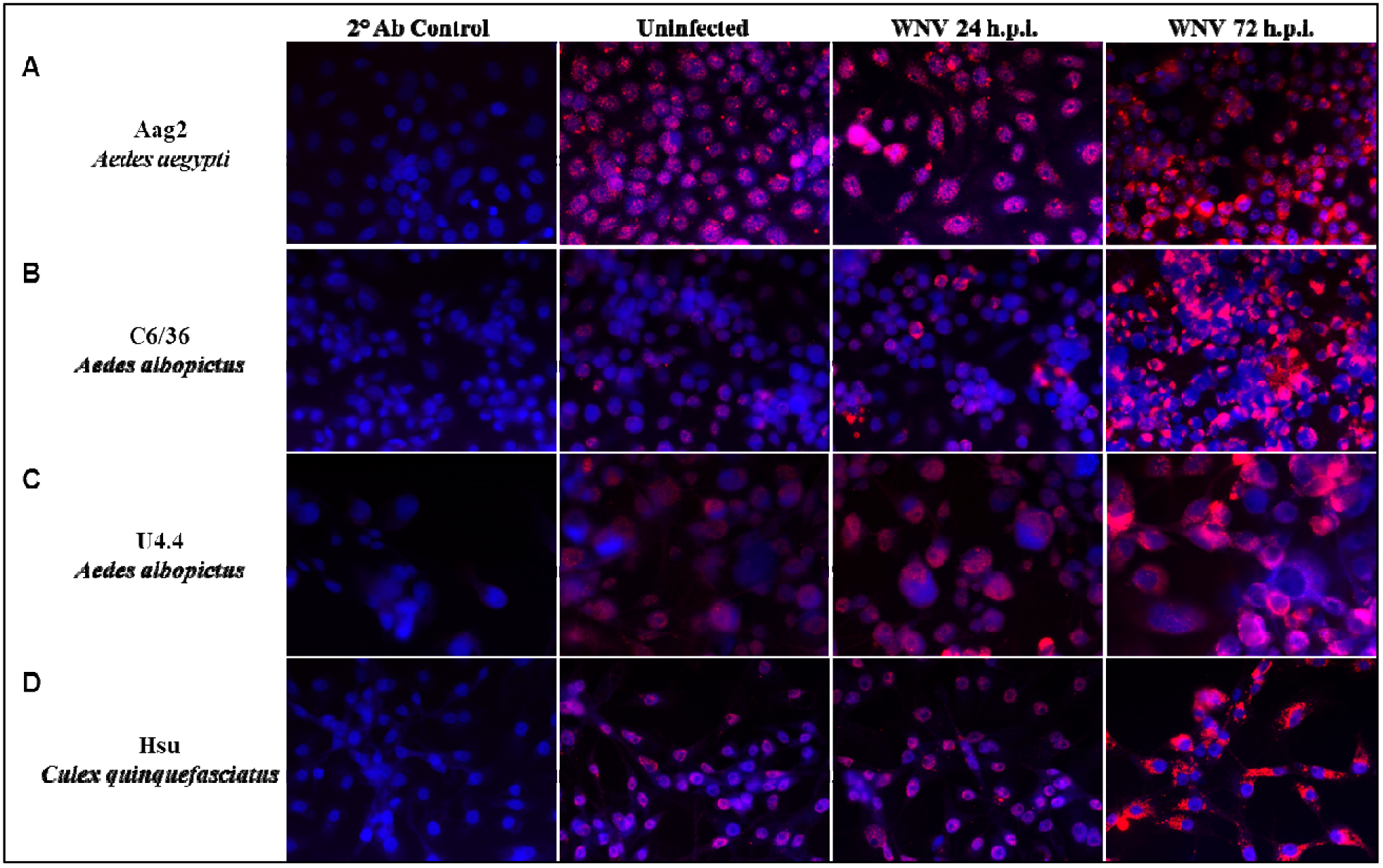
Mosquito cell lines contain dsRNA. Aag2 (A), C6/36 (B), U4.4 (C) and Hsu (D) cells were infected with WNV and fixed at 24 or 72h post infection. Uninfected cells were fixed in parallel. All cells were stained with J2 primary antibody and rabbit anti-mouse Alex-Fluor 555 secondary antibody. A secondary antibody only control (2° Ab Control) is shown to confirm that the observed fluorescence is not due to non-specific binding of the secondary antibody. All images were taken at 63X magnification.

### Viral sequences are abundant in mosquito cell lines

To confirm our immunostaining results indicating virus infection and to identify possible viruses, we implemented an unbiased NGS approach to sequence total RNA from the cell lines. We identified a number of sequences with similarity to virus genomes (Table 2). As expected, we found sequences aligning to PCLV and CFAV in Aag2 cells (Maringer et al., 2017; Schultz et al., 2018; Scott et al., 2010; Stollar and Thomas, 1975). Sequences aligning to PCLV shared 99.8% nucleotide sequence similarity with the Aag2 Bristol strains. We recovered reads spanning 99.9% of each of the three PCLV segments, L, M and S. Similarly the reads aligning to CFAV shared 99.8% sequence similarity to the Aag2 Bristol strain of CFAV with 94.3% of the genome covered. Sequences aligning to WNV and Culiseta flavivirus were also identified; however, we believe these may represent contaminants introduced during library preparation, as libraries from both of these viruses had been recently prepared in the same laboratory (Grubaugh et al., 2016; Misencik et al., 2016). To determine whether the CFAV and PCLV sequences stem from active viral replication, integration into the cellular genome or possibly both, we confirmed the presence of viral RNA and DNA in the cell, as well as RNA in the cell supernatant prior to and following RNase A treatment. PCLV RNA was detected in the cellular lysate and supernatant, however, RNA was not detected after RNase A treatment of the supernatant prior to RNA extraction. Interestingly, we did not detect PCLV in the cellular DNA suggesting that the PCLV sequences are not derived from genomically integrated viral elements. Despite the lack of RNase A protected viral RNA in the supernatant, the almost complete coverage across the length of the genome suggests active replication. CFAV was detected in both DNA and RNA forms in the lysates and supernatants with and without RNase A suggesting that CFAV is actively replicating in Aag2 cells and extrachromosomal or integrated DNA forms exist in the cell.

**Table 2:** Viruses identified by NGS and RT-PCR in mosquito cell lines.

| Cell Line | Closest Virus Match | Accession # | Virus Family | Largest Contig (nt) | % nt identity | % Coverage | Lysate |  | Supernatant |  |
| --- | --- | --- | --- | --- | --- | --- | --- | --- | --- | --- |
|  |  |  |  |  |  |  | RNA | DNA | RNA (-RNase) | RNA (+RNase) |
| Aag2 | Phasi Charoen like virus L segment | KU936055.1 | *Bunyaviridae* | 6789 | 99.8 | 99.9 | + | - | + | - |
|  | M segment | KU936056.1 |  | 3847 | 99.8 | 99.9 | n.d. | n.d. | n.d. | n.d. |
|  | S segment | KU936057.1 |  | 1332 | 99.8 | 99.9 | n.d. | n.d. | n.d. | n.d. |
|  | Cell Fusing Agent virus | KU936054.1 | *Flaviviridae* | 10,067 | 99.8 | 94.3 | + | + | + | + |
| U4.4 | Culex Y Virus segment A | JQ659254.1 | *Birnaviridae* | 2778 | 99.5 | 89.6 | + | + | + | - |
|  | Segment B | JQ659255.1 |  | 2959 | 99.3 | 94 |  |  |  |  |
| Hsu | Merida Virus | KU194360.1 | *Rhabdoviridae* | 11,715 | 95.8 | 99.3 | + | - | + | - |

C6/36 and U4.4 cells are both derived from *Aedes albopictus* and, in fact, are subclones of the original culture (Igarashi, 1978; Miller and Brown, 1992; Singh, 1967). It is unclear when the two cell lines were cultured and maintained separately, but presumably occurred during cloning experiments of Singh’s *Ae. albopictus* (SAAR) cells in 1978 when the C6/36 subclone was first isolated (Igarashi, 1978). While some putative virus-like contigs with partial or disrupted ORFs were identified in each of the three C6/36 cell batches, signatures of bona fide viruses were not identified. In contrast, in U4.4 cells we identified reads aligning to Culex Y virus (CYV), a bisegmented member of the family *Birnaviridae* that was recently identified in *Culex pipiens* mosquitoes (Marklewitz et al., 2012). The fact that CYV was identified in U4.4 cells and not C6/36 cells suggests that CYV infection of U4.4 cells occurred after the two cell lines were subcloned or that infection was lost in one lineage. Reads aligning to segment A had 99.5% nucleotide sequence similarity and covered 89.6% of the segment and reads aligning to segment B had 99.3% similarity spanning 94% of the segment. While we were able to detect RNA and DNA forms of CYV in the lysates and supernatants, we did not detect CYV RNA following RNase A treatment; however, as before, the identification of almost complete genome sequences from both segments would suggest that CYV is actively replicating in U4.4 cells.

We identified one full length genome with nucleotide sequence similarity (95.8%) to Merida virus (MERDV) in the *Cx. quinquefasciatus* Hsu cells. MERDV is a member of the family *Rhabdoviridae* and was initially discovered in pools of *Cx. quinquefasciatus* and *Ochlerotatus* spp. mosquitoes (Charles et al., 2016). MERDV RNA could be detected in both the lysates and supernatants, but could not be detected in cellular DNA or RNase A treated supernatants. Attempts to passage this virus on C6/36 cells were unsuccessful (data not shown). Despite these results, the fact that we identified the near complete genome suggests that MERDV is actively replicating in Hsu cells.

## Discussion

It is well known that insects, including mosquitoes, harbor numerous insect-specific viruses in both wild and laboratory populations (Bolling et al., 2015; Fauver et al., 2016b; Li et al., 2015; Shi et al., 2016). Because mosquito cell lines are generated from these source materials, it is not surprising that ISVs have been identified in mosquito cell cultures (Maringer et al., 2017; Scott et al., 2010). However, the prevalence and taxonomic composition of ISVs in mosquito cell culture systems are unknown. Accordingly, in this study we characterized the viromes of four commonly used mosquito cell lines: Aag2, C6/36, U4.4, and Hsu cells.

From our NGS data we were able to fully reconstruct MERDV; however, we were unable to detect RNase A protected viral RNA in the cell supernatant and, as others have demonstrated, we were unable to recover MERDV after passage on C6/36 cells (Charles et al., 2016). Similarly, despite identifying an almost complete genome from PCLV and, as described by others, the presence of PCLV proteins (Maringer et al., 2017), we were unable to detect protected RNA in Aag2 cell supernatant. Further, we did not detect signatures of either virus in the cellular DNA suggesting that these were not products of integrated sequences. It is possible that these viruses are more prone than other viruses to RNase A degradation. In fact, a previous study found that *Bunyaviridae* nucleocapsids are susceptible to RNase A concentrations of 100 ug/ ml, the concentration used in the preparation of our samples (Osborne and Elliott, 2000). This could explain our inability to detect PCLV in RNase A treated supernatants. It is unclear if similar concentration-dependent RNase A sensitivity could explain our inability to detect encapsidated MERDV which is a member of the family *Rhabdoviridae*. Regardless, the presence of a fully intact MERDV genome and the lack of integration is highly suggestive that MERDV maintains an active and persistent infection of Hsu cells.

Immunofluorescence staining of C6/36 cells revealed the presence of dsRNA and presumably viruses; however, our NGS analysis did not identify signatures of bona fide viruses. These incongruent findings suggest that intracellular dsRNA originate from sources other than actively replicating virus such as cellular dsRNA molecules. We did identify a number of partial or disrupted virus-like ORFs and it is therefore possible that the dsRNA signal could arise from endogenous viral elements (EVE). Mechanistically this could occur by EVE transcripts folding back upon themselves or from convergent EVE transcripts, both of which would generate dsRNA substrates. The origin of the EVEs is unknown, but likely represent ancient viral infections that had integrated into the genome of the original cell culture source.

C6/36 cells lack a functional antiviral RNAi pathway (Brackney et al., 2010) which makes them ideal for propagating arboviruses in the laboratory. This deficiency has been mapped to a homozygous frame-shift mutation in the *dcr-2* gene resulting in a premature termination codon (Morazzani et al., 2012). Conversely, U4.4, Aag2 and Hsu cells have fully functional RNAi responses and, consequently, propagation of viruses in these cell lines typically generates much lower titers than C6/36 cells (Paradkar et al., 2012; Scott et al., 2010; Siu et al., 2011). In this study we demonstrate that U4.4, Aag2, and Hsu cells are persistently infected with viruses, but that C6/36 cells have remained uninfected. The symmetry between viral infection and RNAi functionality is intriguing. Based on our findings, we speculate that because C6/36 cells are maintained in a sterile culture environment devoid of viruses there is no selective advantage to maintain a functional antiviral RNAi pathway. Consequently, once a mutation arose in the *dcr-2* gene it was able to spread throughout the culture without the culture experiencing fitness losses.

Numerous endogenous flavivirus elements have been identified in mosquito cells and mosquitoes suggesting that genomic incorporation of RNA viruses is a common occurrence (Crochu et al., 2004; Suzuki et al., 2017). The process by which the genomes of non-retrovirus RNA viruses can integrate was recently described by Goic *et al.* (2013 & 2016). They found that the integration of RNA virus genomes into the genomes of mosquito cells and mosquitoes is mediated by endogenous retrotransposon reverse transcriptase activity. It is thought that this process helps control persistent RNA viruses. All of the viruses reported here are RNA viruses, yet we detected DNA forms of CFAV and CYV and not PCLV or MERDV. This could represent a discrepancy in the template selection preferences associated with this process. In fact, this endogenous retrotransposon reverse transcriptase mediated process has only been demonstrated for positive-sense single-stranded RNA viruses like CFAV (Goic et al., 2016; Goic et al., 2013) and it is therefore not surprising that DNA forms of CFAV were detected. Interestingly, CYV is a member of the family *Birnaviridae* and has a double stranded RNA genome. The presence of CYV DNA forms suggests that dsRNA can also serve as a template for the production of extrachromosomal or integrated viral DNA; however, we were unable to detect DNA forms of either PCLV (*Bunyaviridae*) or MERDV (*Rhabdoviridae*) both of which have negative-sense single-stranded RNA genomes. Genetic signatures of other negative strand virus have been found integrated into the *Aedes aegypti* genome (Katzourakis and Gifford, 2010) which suggests that genetic elements derived from PCLV and MERDV have not yet integrated into the genome or that integrated elements exist but have diverged from the consensus sequence and, therefore, were not efficiently amplified during PCR.

It is unclear at this time how or if these persistent viral contaminants have affected the outcomes of mosquito cell-based studies. It is known that viruses can drastically alter cellular homeostasis, lipid levels and distribution, organelle abundance and integrity, cellular RNA levels, protein abundance and antiviral defense mechanisms. For example, members of the family *Bunyaviridae*, like PCLV identified in Aag2 and C6/36 cells, drastically alter cellular mRNA levels through “cap-snatching” and viral suppressors of RNA interference have been identified in numerous insect-specific viruses (Hopkins et al., 2013; van Cleef et al., 2011). In addition, it is known that viruses can interact with one another. Recently, it was demonstrated that CFAV and dengue virus can synergistically promote the replication of one another during infection of Aag2 cells (Zhang et al., 2017). Conversely, others have found that these viruses can interfere with arboviral replication (Goenaga et al., 2015). Such findings suggest that the persistent viral infections identified in this study could affect the outcomes of arbovirus evolution and virus-vector interaction studies performed in these cell lines. Clearly defining model systems is paramount to properly interpreting results generated in these systems. This study better defines mosquito cell culture systems, the results from which can be used to improve experimental design and interpretation of results.

## Acknowledgements

This work was supported in part by grants from the Centers for Disease Control and Prevention (U50/CCU116806-01 and U01/CK000509-01), the US Department of Agriculture Hatch Funds and Multistate Research Project (CONH00773 and NE1443), and the National Institute of Health, National Institute of Allergy and Infectious Diseases (AI067380), and NIH/NCATS (UL1 TR001082).

## Contributions

Conception and design: J.W-L., C.R., N.D.G., P.M.A., D.E.B, G.D.E. Acquisition of data: J.W-L., C.R., M.J.M. and N.D.G. Analysis and interpretation of data: M.D.S., J.W-L., and C.R. Writing of the manuscript: D.E.B, P.M.A., J.W-L. and C.R. Study supervision: P.M.A., D.E.B and G.D.E. All authors have critically evaluated and approved the final manuscript.

